# Resurrecting self-cleaving mini-ribozymes from 40-million-year-old LINE-1 elements in human genome

**DOI:** 10.1101/2021.04.06.438727

**Authors:** Zhe Zhang, Peng Xiong, Junfeng Wang, Jian Zhan, Yaoqi Zhou

## Abstract

Long Interspersed Nuclear Element (LINE) retrotransposons play an important role in genomic innovation as well as genomic instability in many eukaryotes including human. Random insertions and extinction through mutational inactivation make them perfectly time-stamped “DNA fossils”. Here, we investigated the origin of a self-cleaving ribozyme in 5’ UTR of LINE-1. We showed that this ribozyme only requires 35 nucleotides for self-cleavage with a simple but previously unknown secondary-structure motif that was determined by deep mutational scanning and covariation analysis. Structure-based homology search revealed the existence of this mini-ribozyme in anthropoids but not in prosimians. In human, the most homologs of this mini-ribozyme were found in lineage L1PA6-10 but essential none in more recent L1PA1-2 or more ancient L1PA13-15. We resurrected mini-ribozymes according to consensus sequences and confirmed that mini-ribozymes were active in L1PA10 and L1PA8 but not in L1PA7 and more recent lineages. The result paints a consistent picture for the emergence of the active ribozyme around 40 million years ago, just before the divergence of the new world monkeys (Platyrrhini) and old-world monkeys (Catarrhini). The ribozyme, however, subsequently went extinct after L1PA7 emerged around 30 million years ago with a deleterious mutation. This work uncovers the rise and fall of the mini-LINE-1 ribozyme recorded in the “DNA fossils” of our own genome. More importantly, this ancient, naturally trans-cleaving ribozyme (after removing the non-functional stem loop) may find its modern usage in bioengineering and RNA-targeting therapeutics.

## Introduction

Ribozymes (catalytic RNAs) are thought to dominate early life forms (1) until they were gradually replaced by more effective and stable proteins. One of the few that remain active is self-cleaving ribozymes found in a wide range of living organisms (2), involving in rolling circle replication of RNA genomes (3–5), biogenesis of mRNA and circRNA, gene regulation (6–9), and co-transcriptional scission of retrotransposons (10, 11). Involving in mobile retrotransposons indicates the importance of self-cleaving ribozymes in the overall size, structure, function and innovation of the genomes.

Most retrotransposon-related self-cleaving ribozymes found so far are observed in non-mammalian genomes. They belong to the two most widespread ribozyme families: hepatitis delta virus (HDV)-like ribozymes in R2 elements of *Drosophila simulans* (10) and L1Tc retrotransposon of *Trypanosoma cruzi* (11) and hammerhead ribozymes (HHR) in short interspersed nuclear elements (SINEs) of *Schistosomes* (12), Penelope-like elements (PLEs) of different eukaryotes (13, 14) and retrozymes in plants (15). The only exception is the Long Interspersed Nuclear Element-1 (LINE-1) ribozyme located inside the 5′ untranslated region (UTR) of a LINE-1 retrotransposon. It was discovered along with the CPEB3 ribozyme in a biochemical selection-based experiment (8). Unlike CPEB3 ribozyme, there was no follow-up study. As a result, little is known about the LINE-1 ribozyme.

In this study, we investigate the secondary structure motif and minimal function region of the LINE-1 ribozyme by deep mutational scanning and covariation analysis (16). The secondary structure obtained allows us to perform the structure-based homolog search against the reference genome and experimentally test the activity of the consensus sequences in subfamilies of LINE-1. We found that the LINE-1 ribozymes belong to a lineage of LINE-1 which was active approximately 40 million years ago, and are widespread in most primate genomes.

## Result

### Deep mutational scanning of LINE-1 ribozyme

Our previous work indicates that deep mutational scanning can lead to co-variation signals for highly accurate inference of base-pairing structures (16). Here, the same method is applied to the full-length LINE-1 ribozyme (146 nucleotides, LINE-1-fl). In this method, high-throughput sequencing can directly measure the relative cleavage activity (RA) of each variant in a library of LINE-1-fl RNA mutants according to the ratio of cleaved to uncleaved sequence reads of the mutant, relative to the same ratio of the wild-type sequence. Figure 1a shows the average RA for a given sequence position for all the mutants with a mutation at the position. The activity of the LINE-1 ribozyme is only sensitive to the mutations in two short segments (54-71 and 83-99) where the RA can be severely reduced. Thus, the two terminal ends and the central region between 72 and 82 are not that important for the self-cleaving activity.

**Figure 1.**
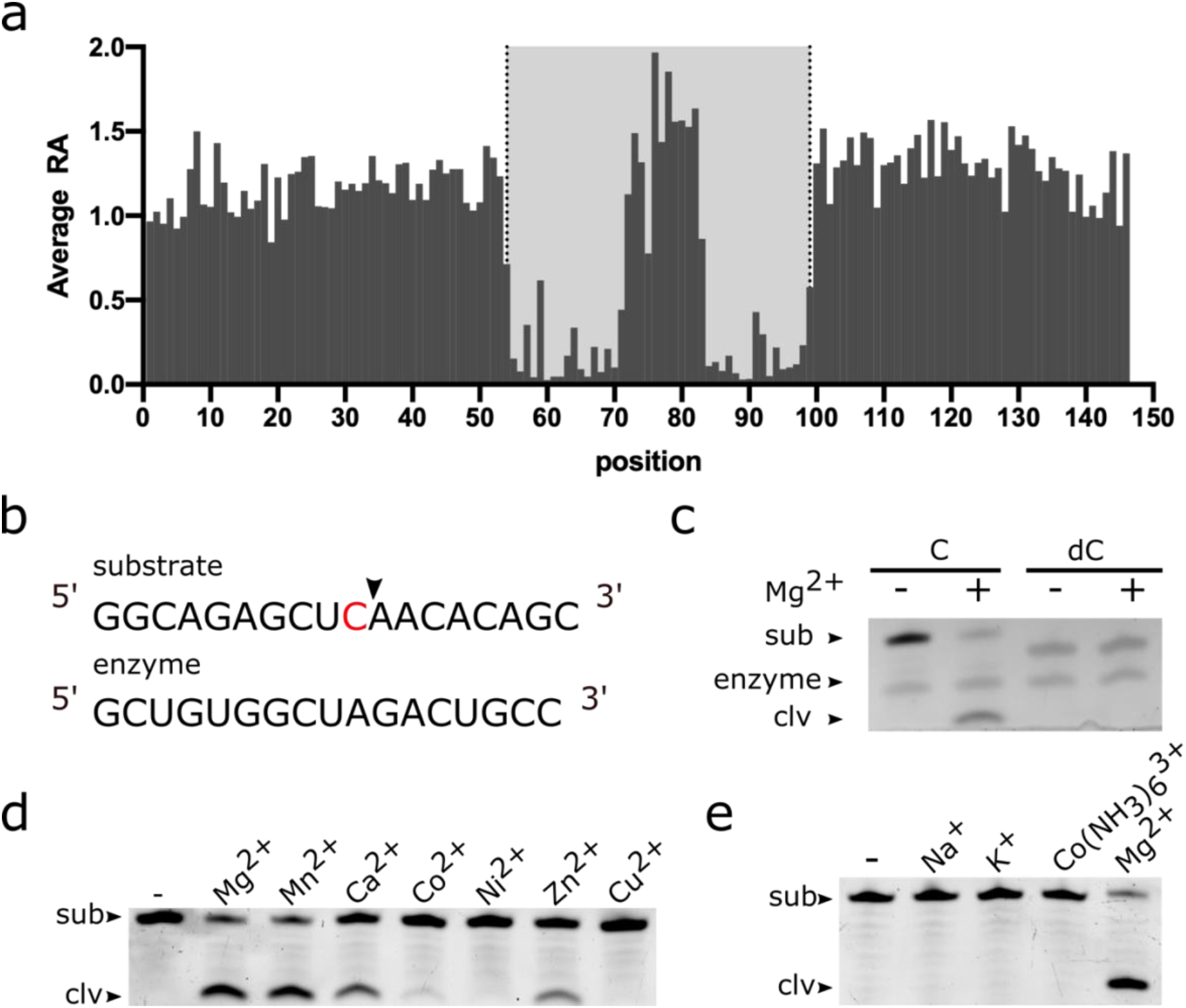
(a) Average relative activity of mutations at each sequence position of the full-length LINE-1 ribozyme (LINE-1-fl). (b) The components of the bimolecular construct of the minimal ribozyme, substrate strand (54-71) and enzyme strand (83-99). The arrowhead indicates the cleavage site. (c) PAGE-based gel cleavage assays of the bimolecular construct with the original substrate (C) and a substrate analog (dC) wherein a 2′-deoxycytosine is substituted for the cytosine ribonucleotide at position 10 of the substrate RNA. The substrate RNA was incubated with the enzyme RNA at 37°C for 1h in the presence (+) or absence (−) of 1mM MgCl_2_. (d) Cleavage assays in the absence (−) or presence (+) of various divalent metal ions at a concentration of 1mM. (e) Cleavage assays in the absence (−) or presence of monovalent cations, cobalt hexamine chloride [Co(NH_3_)_6_Cl_3_] or MgCl_2_ at 5mM for 1h.

### Activity confirmation and biochemical analysis of LINE-1-mini ribozyme

To confirm the cleavage activity of the minimal region, we obtained the segment 54-71 as the substrate strand with the cleavage site and the segment 83-99 as the enzyme strand (Figure 1b). When the substrate strand is mixed with the enzyme strand, the majority of the substrate strand was cleaved in the presence of Mg^2+^ (Figure 1c, 1d, 1e). This construct contains 35 nucleotides in total, which makes it the shortest self-cleaving ribozyme reported so far. The observed rate constant (k_obs_) value for this construct was 0.04254 min^−1^ with 1mM Mg^2+^ and pH 7.5. In all the previously reported self-cleaving ribozyme families, the self-cleaving reaction occurs through an internal phosphoester transfer mechanism that the 2′-hydroxyl group of the −1 (relative to the cleavage site) nucleotide attacks the adjacent phosphorus resulting in the release of the 5′ oxygen of +1 (relative to the cleavage site) nucleotide (2, 17–20). We confirmed that the LINE-1-mini ribozyme uses the same mechanism because an analog RNA of LINE-1 ribozyme that lacks the 2′ oxygen atom in the −1 nucleotide (dC10) is unable to cleave (Figure 1c). We further examined the dependence of the metal ions on the cleavage activity. At a concentration of 1mM, ribozyme cleavage can be observed with Mg^2+^, Mn^2+^, Ca^2+^ and Zn^2+^ but little or none with Co^2+^, Ni^2+^, Cu^2+^, Na^+^, K^+^ or Co(NH_3_)_6_^3+^, indicating that direct participation of specific divalent metal ions is required for self-cleavage.

### Deep mutational scanning of LINE-1-mini ribozyme

We employed the contiguous minimal functional segment (54-99 or 46 nucleotides, LINE-1-mini) for the second round of deep mutational scanning. Performing the second round is necessary because the deep mutational scanning of the full-length LINE-1 ribozyme has a low 18.5% coverage of double mutations (Supplementary Table S1) that is not sufficient for high-resolution inference of secondary structure. This second round employed a chemically synthesized doping library with a doping rate of 6%, rather than a mutant library generated from error-prone PCR that was biased toward the sequence positions with A/T nucleotide (Supplementary Table S2 and Figures S1 and S2). In addition, to amplify the signals of cleaved RNAs after *in vitro* transcription of the mutant library of LINE-1-mini, we selectively capture cleaved RNAs by employing RtcB ligase and a 5′-desbiotin, 3′-phosphate modified linker because they only react with the 5′-hydroxyl termini that exist only in cleaved RNAs (21). The captured active mutants were further enriched by the streptavidin-based selection after ligation. The technology improvement leads to less biased mutations in terms of positions (Supplementary Figure S1) and mutation types (Supplementary Figure S2). More importantly, it achieves 99.3% and 99.9% coverage of single and double mutations, respectively, within the contiguous minimal functional segment (Supplementary Table S1).

### Inference of secondary structure by covariation-induced deviation from activity (CODA)

We performed the CODA analysis (16) based on the relative activities of 45,925 and 72,875 mutation variants obtained for the full length and minimal LINE-1 ribozyme, respectively. CODA detects base pairs by locating the pairs with large covariation-induced deviation of the activity of a double mutant from an independent-mutation model. Figure 2a shows that both full-length and minimal ribozyme data reveal the existence of two stem regions. The latter, in particular, clearly shows two stems with lengths of 6 nt and 5 nt, respectively, as a result of nearly 100% coverage of single and double mutations for the minimal region. Moreover, it suggests a non-canonical pair 6A41A at the end of the first stem, although the signal is weak. A weak signal is expected because non-canonical pair is stabilized by the weak tertiary interaction only. There are some additional weak signals in the LINE-1-mini result. Most of them are a few sequence distance apart (|i-j|<6, local base pairs). We found that some of them are caused by a high relative activity (RA’, measured in the second round of mutational scanning) of the double mutant and not likely to be true positives because the corresponding single mutations are not very disruptive (RA’ > 0.5).

**Figure 2.**
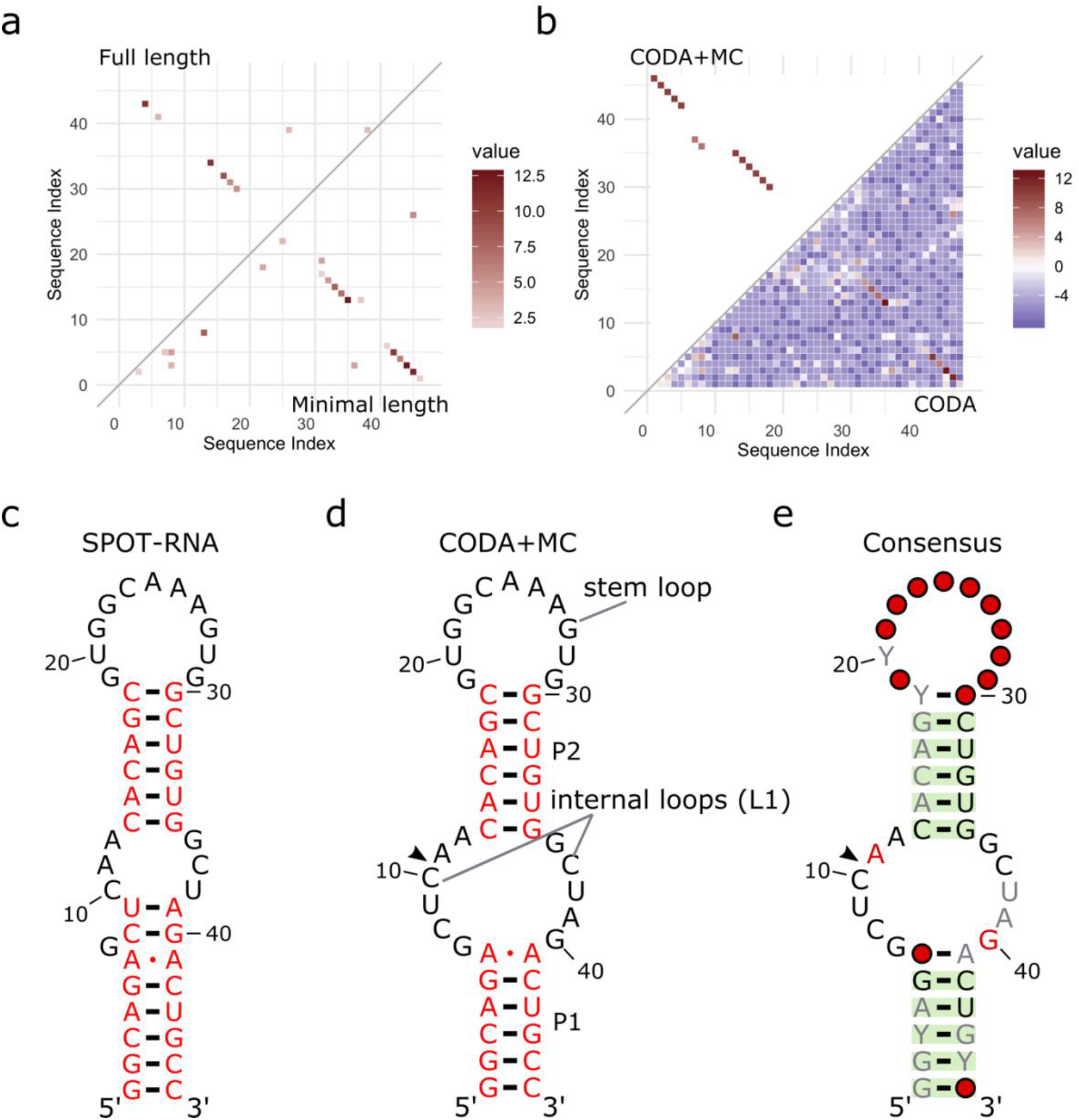
(a) Comparison between the base pairing map (probability score value > 2) inferred from covariation-induced-deviation-of-activity (CODA) analysis of deep mutational scanning of the minimal contiguous LINE-1 ribozyme (LINE-1-mini) (lower triangle) and that from the corresponding region (54-99 nt) in the deep mutational scanning of the full-length LINE-1 ribozyme. (b) Comparison between the base pairing map inferred from LINE-1-mini deep mutational data by CODA (lower triangle) and that after Monte-Carlo simulated annealing (CODA+MC, upper triangle). (c) Secondary structure model of LINE-1-mini ribozyme predicted by the SPOT-RNA secondary structure predictor. (d) Secondary structure model inferred from the deep mutational scanning result. (e) Consensus sequence and secondary structure model for LINE-1-mini ribozyme based on the alignment of 1394 functional mutants. The arrowhead indicates the ribozyme cleavage site. Positions with conservation of 95, 98, and 100% were marked with gray, black and red nucleotide, respectively; positions in which nucleotide identity is less conserved are represented by circles. Green shading denotes predicted base pairs supported by covariation. Y denotes pyrimidine.

### Secondary structure by Monte Carlo simulated annealing and SPOT-RNA

Probabilistic CODA results were combined with experimental pairing energy functions for global minimum search by Monte Carlo simulated annealing (CODA+MC) as in the previous method (16). The resulting base pairs are shown in Figure 2b (upper triangle). Those mostly local false positives disappeared. However, the noncanonical AA pair was disappeared as well, likely because the energy function was not optimized for noncanonical pairs. A new 18C30G pair added is a natural extension of the second stem. Two separate consecutive base pairs (7G37C, 8C36G) with relatively low signals were also added in the CODA+MC result. These two base pairs are likely false positives because they do not appear in all the models generated from Monte Carlo simulated annealing with a low probability of 0.52. As a comparison, we also predicted the secondary structure for LINE-1-mini by a recently developed SPOT-RNA (22) (Figure 2c). The predicted structure has exactly the same two stems from the MC simulation as well as a non-canonical pair 6A41A. However, it has two extra base pairs (8C40G and 9U41A) that are not supported by co-mutational signals (Figures 2a and 2b). It is interesting, however, that they were consistent with the homology model as shown below. Taking all together leads to a confident secondary structure for the LINE1-mini ribozyme, which contains two stem regions (P1, P2), two internal loops and a stem loop (Figure 2d).

### Consensus sequence of LINE-1-mini ribozyme

The consensus sequence for LINE-1-mini ribozyme was created by R2R (23) by aligning 1394 mutants with RA ≥ 0.5. Figure 2e overlays the consensus sequence onto the secondary structure model from the CODA+MC analysis. The self-cleavage of the LINE-1-mini occurs between a conserved CA dinucleotide which locates inside the upstream part of the internal loops (L1) linking P1 and P2. The stem loop region showed the lowest sequence identity, consistent with its insensitivity to mutations in Figure 1a, providing additional support for the secondary structure model obtained here.

### Homology modelling of LINE-1-mini ribozyme

We found that the above two internal loops with 5 and 6 nucleotides, respectively, differ from the catalytic internal loops of twister sister only by one base in each loop (Figure 3a) (24). The conservation of the cleavage site along with the surrounding internal loops suggests the underlying similar cleavage mechanism and three-dimensional structure. Figure 3b shows the homology modelling structure of LINE-1-mini ribozyme (internal loops plus stems) by simply copying the backbone and base conformations from twister sister to LINE-1-mini ribozyme along with an extension of the stem regions. The structure reveals the internal loops L1 as the catalytic region involving a guanine–scissile phosphate interaction (G25–C10-A11), a splayed-apart conformation of the cleavage site, additional pairings and hydrated divalent Mg^2+^ ions. As shown in Figure 3c-f, stem P1 is extended by the Watson-Crick U9A28 which forms part of G7 (U9A28) base triple and Watson-Crick C8G29, whereas stem P2 is extended through trans non-canonical A12G25 and trans sugar edge-Hoogsteen A11C26. While the cleavage site C10-A11 is splayed apart, with C10 directed outwards and A11 directed inwards into the ribozyme fold. A11 (anti alignment) is anchored in place by stacking between Watson-Crick U9A28 and non-canonical trans A12G25 pairs and hydrogen bond between N7 atom of A11 and O2’ atom of C26. The non-bridging phosphate oxygens of the C10A11 scissile phosphate form hydrogen bonds to the N1 of G25. Moreover, all the additional base pairs in L1 are relatively conserved, as new base pairs formed after mutations showed RA < 0.5 according to deep mutational scanning result. This is consistent with the stacking, long-range pairing and hydrogen-bonding interactions in the L1 region.

**Figure 3.**
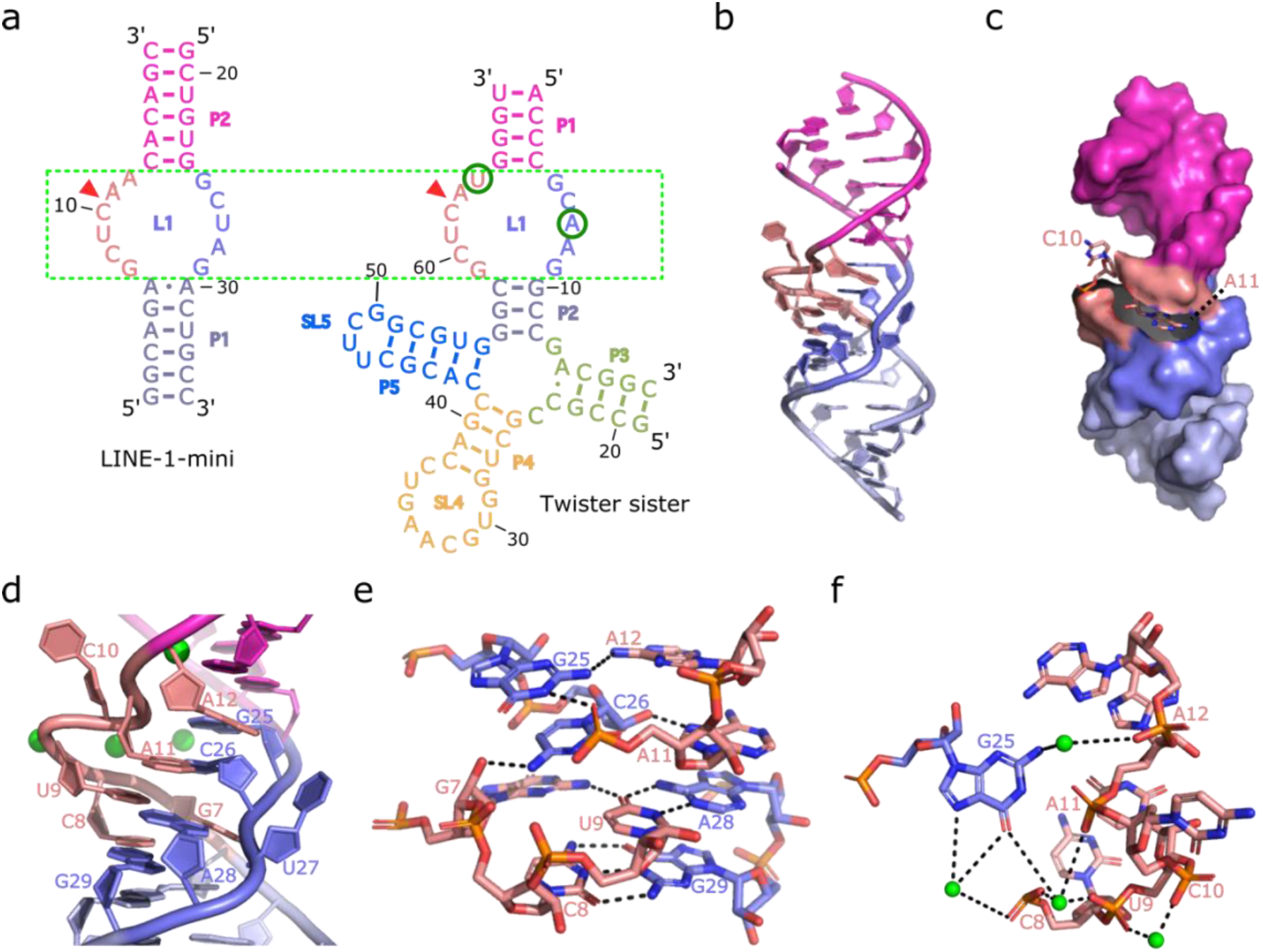
(a) Comparison between the secondary structure of the bimolecular construct of LINE-1-mini (left) and that of the four-way junctional twister-sister ribozyme (right, PDB ID: 5Y87) reveals strikingly similar internal loops surrounding the cleavage site (red triangles) with a single base difference at each side of the internal loops L1. (b) A ribbon view of the homology modelling structure of the LINE-1-mini color-coded as shown in (a) left. (c) The homology modelling structure of the LINE-1-mini ribozyme with the RNA in a surface representation, except for the cleavage site C10-A11, which is shown in a stick representation. (d) A detailed view of the catalytic center. The divalent metal ions in the tertiary structure are shown as green balls. Bases C10 and A11 at the cleavage site are splayed apart, with C10 directed outwards and A11 directed inwards into the ribozyme fold. A11 (anti alignment) is anchored in place by stacking between Watson-Crick U9A28 and non-canonical trans A12G25 pairs. (e) Additional interactions found in the homology modelling structure of the LINE-1-mini ribozyme. Stem P1 is extended by the Watson-Crick U9A28 which forms part of G7(U9A28) base triple and Watson-Crick C8G29, whereas stem P2 is extended through trans non-canonical A12G25 and trans sugar edge-Hoogsteen A11C26. The non-bridging phosphate oxygens of the C10A11 scissile phosphate form hydrogen bonds to the N1H of G25. (f) Hydrogen bond networks in internal loops L1 involving a set of four divalent Mg^2+^ ion.

### Homology search by BLAST-N

To locate close homologs, we performed BLAST-N search (25) in the NCBI RefSeq Genome Database using the sequence of LINE-1-mini. 905 homologs were found in primate genomes only, with 11-58 homologs per primate genome (Supplementary Figure S3). However, there was no exact match. Only one has a single mutation (U38C) in the genome assembly of Nomascus leucogenys (nomLeu3). The majority of these homologs have 3 or more mismatches (Supplementary Figure S4). The sequence with the highest sequence identity in human genome assembly (hg38) has 3 mismatches but is located in a nonactive LINE-1 subfamily (see below).

### Structure-based homology search by Infernal

To locate more homologies, we performed cmsearch based on a covariance model (26) built on the sequence and secondary structural profiles because most sequence homologs above maintain the base pairings in the stem regions (Supplementary Figure S5). Supplementary Table S3 listed all cmsearch results from 18 representative primate genomes. The majority (15 out of 18) of primate genomes have a large number of LINE-1 homologs (>500) and the remaining three have essentially none. Interestingly, almost all search results were mapped to the LINE-1 retrotransposon according to the RepeatMasker annotation. Supplementary Figure S6 further shows that these LINE-1 ribozymes are conserved inside the 5′ UTR, located inside the 650-750 nt region upstream of the ORF. Despite of using different sequence databases, BLAST-N and Infernal both located the closest single (single mutation of U38C) in the genome of Nomascus leucogenys (Gibbon).

### Evolution of LINE-1-mini ribozyme in primate species

The existence of LINE-1 ribozymes in anthropoids such as human, chimpanzee and bonobo but not in prosimians (tarsier, mouse lemur and bushbaby) (Supplementary Table S3) is consistent with the evolution of primate species. As shown in the phylogenetic tree (http://genome.ucsc.edu) in Figure 4a, the lineage of ribozyme-containing LINE-1 was generated just before the divergence of the new world monkeys (Platyrrhini) and old-world monkeys (Catarrhini), which was approximately more than 40 million years ago.

**Figure 4.**
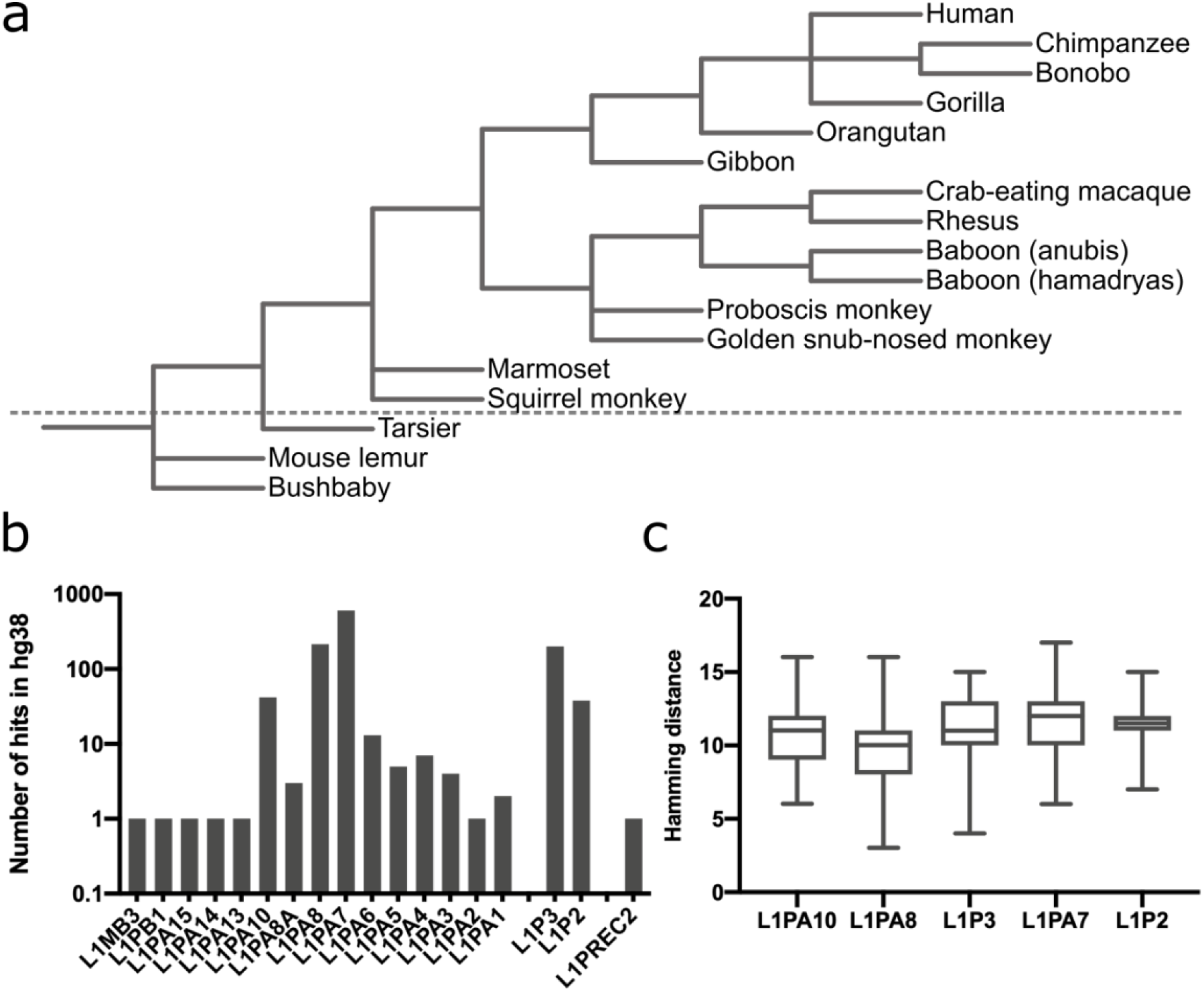
(a) Phylogenetic tree of representative primate species. The species showed above the dotted line contains around 1000 homologs of LINE-1-mini, while the species under the dotted line do not contain LINE-1-mini homolog. (b) The distribution of LINE-1-mini homologs in different LINE-1 subfamilies of human genome assembly (hg38). L1PA, L1PB and L1MB subfamilies are classified based on 3′ UTR, L1P subfamilies are classified based on ORF2. L1PA8A is a side branch from the L1PA8. L1PA1 is also known as L1Hs, and L1P3 is equal to L1PA7-9, L1P2 is equal to L1PA4-6. (c) The hamming distance distribution of the top 5 abundant LINE-1 subfamilies. The hamming distance is calculated between each homolog and the query sequence.

### Evolution of LINE-1-mini ribozyme in human

We classify the LINE-1-mini homologs found in human genome according to their locations in different subfamilies of LINE-1. These LINE1 subfamilies were classified based on the consensus sequences of the 3′ UTR or ORF2 by using the RepeatMasker classification (27) and named according to their ages (larger number, more ancient if started with the same alphabet). Figure 4b shows the number of LINE-1-mini homologs for those lineages with clear time-stamps (27–29). The most mini-ribozyme homologs were found in lineage L1PA6-10 but essential none (only 1 or 2) in more ancient L1PA13-15 and more recent L1PA2 and L1PA1 (L1Hs). A few found in L1PA13-15, L1PA1-2 and others are likely false positive matches, given the nature of self-duplicating retrotransposons. In Figure 4c we compare the hamming distance of the 5 most abundant subfamilies to the reference sequence. The L1PA8 subfamily is the one with the highest average similarity (least divergent) to LINE-1-mini reference sequence. It also contains the one with the highest identity (3 mismatches) to LINE-1-mini. Note that L1PA8 is active around 40 million years ago (28), consistent with the time for the divergence of the new world monkeys and old-world monkeys.

### Resurrecting self-cleaving mini-ribozymes by consensus

To examine if LINE-1-mini was active when the LINE1 family was current, we reconstructed the putative ancestral representatives of each LINE-1 subfamily according to the majority-rule consensus sequence (30). This is necessary because the current sequence has accumulated mutations up to now. We only chose the five most abundant subfamilies for constructing the consensus sequence because a quality consensus sequence requires enough sequences for extracting statistical information. In a consensus sequence, the nucleotide at each position was determined as the nucleotide with the highest occurrence. Among these 5 subfamilies, the L1PA10 is the most ancient and the L1P2 (L1PA4-6) is the youngest. The consensus sequences of 5 subfamilies of LINE-1 mini ribozyme obtained are shown in Figure 5a.

**Figure 5.**
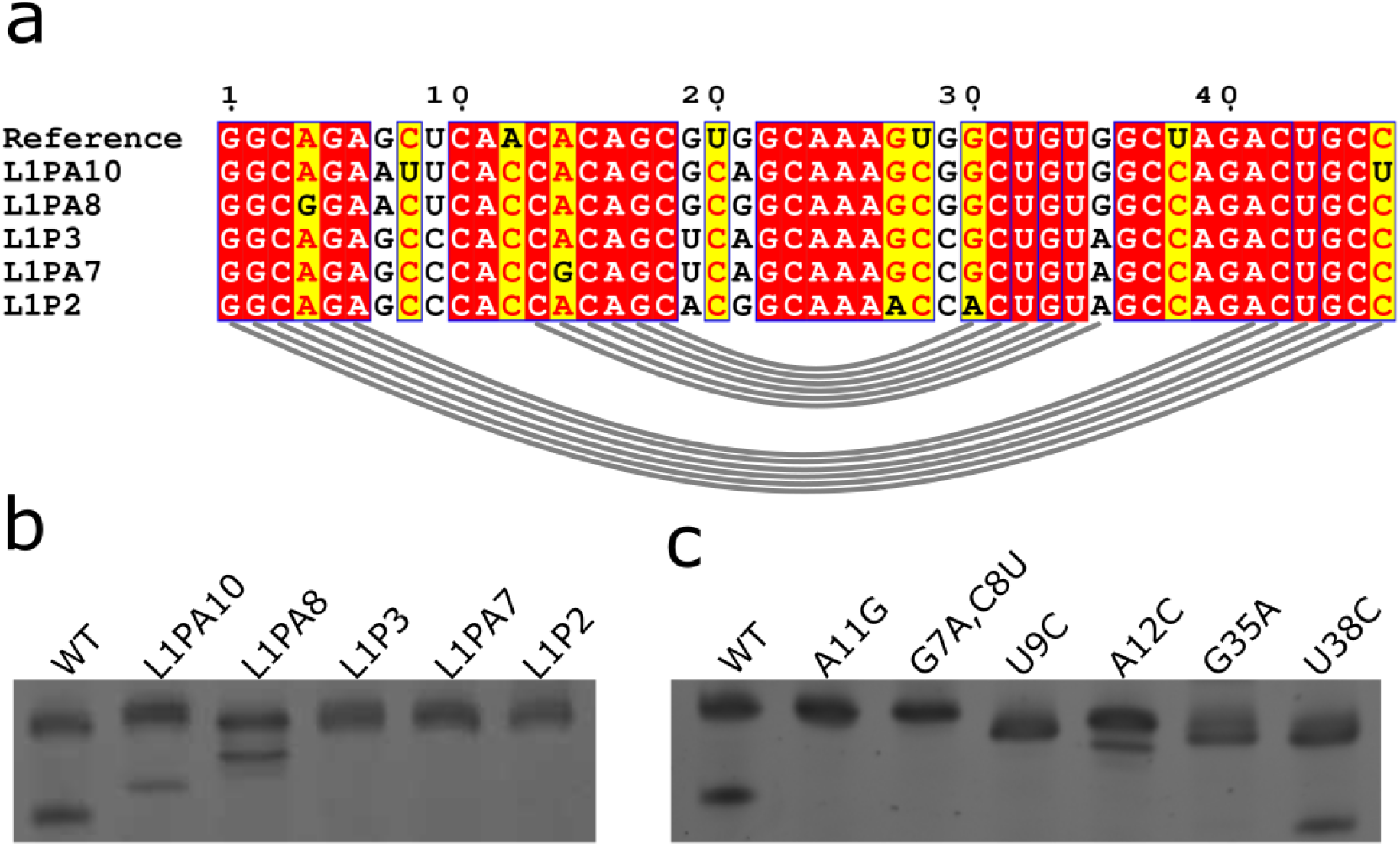
(a) Alignment of consensus sequences from different LINE-1 subfamilies with the reference sequence. (b) PAGE result of *in vitro* transcribed RNA of different consensus sequences. (c) PAGE result of *in vitro* transcribed RNA of different LINE-1 ribozyme mutants.

### Self-cleavage activity of consensus sequences

We tested the cleavage activity of these consensus sequences by PAGE-based gel separation. As shown in Figure 5b, the consensus sequences of LINE1-mini in L1PA10 and L1PA8 are active but not those in L1P3, L1PA7 and L1P2. Unlike the L1PA subfamilies, L1P3 and L1P2 are classified based on ORF2 instead of 3′ UTR. L1P3 is equal to L1PA7-9, and L1P2 is equal to L1PA4-6 (27). What lead to the loss of activity for recent LINE-1 subfamilies but not the earlier ones, given all having least 6 mismatches when comparing with the reference sequence? The reasons are not likely from the two stem regions alone. L1PA10 has C46U, which has a minimal effect because this mutation leads to G1U46 pair. A wobble GU pair is also energetically favourable (31). L1PA8 has one mutation (A4G), which also leads to a G4U43 pair. On the other hand, L1P3, L1PA7 and L1P2 all have a mutation G35A, which disrupted the original C13G35 pair and lead to the loss of activity (Figure 5c). However, a C13U mutation does exist in each subfamily (Supplementary Figure S7), with high frequency in L1P3 (0.25), L1PA7 (0.38) and L1P2 (0.37). This C13U mutation would restore the disruption caused by G35A via forming a new U13A35 pair. Thus, an additional reason must exist for loss of activity in L1P3, L1PA7 and L1P2.

### Deleterious mutations in internal loops

If not in stem regions, the culprit must be in the internal loops as internal loop regions contribute significantly to the self-cleaving activity (Figure 1a). The major changes of the five consensus sequences are in the first internal loop. A12C in all the 5 sequences leads to a one-base pair extension of the second stem (P2) but almost no activity from the deep mutational scanning result (RA=0.03). However, a cleaved band with minor shift could be observed in PAGE separation (Figure 5c). This activity was missed by deep mutational scanning because RA is calculated by counting only the reads matched to the same cleavage site. One common mutation in L1PA7, L1P3 and L1P2 but not in L1PA10 and L1PA8 is U9C and Figure 5c shows that U9C alone can kill the activity. For L1PA10 and L1PA8, the combination of A12C with G7A, C8U or G7A only altered the cleavage site of the mini ribozyme. In the second internal loop, a U38C mismatch is highly conserved in all homologs, which is also found in the homolog sequence with the highest identity in *Nomascus leucogenys* genome. The U38C single mutation has a RA of 0.6 according to the deep mutational scanning result. The *in vitro* cleavage activity of this mutation was further confirmed by using PAGE-based gel separation (Figure 5c). Thus, U9C is likely what kills the activity of recent LINE-1 mini-ribozyme.

## DISCUSSION

The 5′ UTR of LINE-1 retrotransposon is important for the transcription initiation of LINE-1 retrotransposons. Thus, the discovery of a LINE-1 self-cleaving ribozyme (8) poses a question regarding its functional role. However, lacking its structural information makes subsequent studies difficult. In this paper, we applied a recently developed the CODA method (16) to infer the minimal functional region (54-70 and 83-99) from the first round of deep mutational scanning of the full-length ribozyme. This is followed by deep mutational scanning of the minimal region (54-99) to infer its secondary structure. We are confident about the secondary structure obtained because of the consistency between the results from the deep mutational study of the full-length and those from the minimal length of LINE-1 ribozyme. The structure is also supported by secondary structure predictor SPOT-RNA, the recent accurate predictor trained by deep transfer learning (22). Moreover, we found that the internal loop region of LINE-1-mini shared high similarity with the known catalytic region of the twister sister. The conserved cleavage site indicates that they should have a similar structure and mechanism. According to the deep mutational scanning result, two mismatches in the catalytic region A12U and U27A have RA of 0.28 and 0.47, respectively, whereas the coexistence of these two mismatches only has a RA of 0.09. However, under a similar experimental condition, the observed rate constant (k_obs_) for twister sister ribozyme was ~5 min^−1^(20), whereas the k_obs_ for LINE-1-mini was much lower (0.04254 min^−1^). This suggests that long range interactions involving the SL4 region in twister sister ribozyme must have stabilized the catalytic region for the improved cleavage rate.

Armed with the minimal active sequence and confident secondary structure, we searched their homologs by BLAST-N (25) and cmsearch (26). To our surprise, the wide-type sequence does not exist in human genome assembly (hg38) but has a minimum of three mutations which is located in an extinct LINE-1 subfamily (L1PA8). Searching against representative genomes, we found that the genome of *Nomascus leucogenys* is the only one with a single mutation to the wild-type sequence (U38C), which has the in vitro self-cleavage activity (Figure 5c). Moreover, LINE-1 mini ribozyme only exists in anthropoids but not in prosimians, indicating this lineage of ribozyme-containing LINE-1 was generated just before the divergence of the new world monkeys (Platyrrhini) and old-world monkeys (Catarrhini), which was approximately more than 40 million years ago (28, 32). Indeed, with >500,000 copies of LINE-1 in human genomes, we can only find LINE-1 mini ribozymes in LINE-1 subfamilies which were active 40 million years ago. Resurrected mini-ribozymes in different subfamilies confirmed their cleavage activity 40 million years ago but subsequently extinct likely due to a deleterious mutation (U9C).

While the *in vivo* function of this mini-ribozyme may be only historically important in human for regulating the amplification of this LINE-1 lineage, its resurrection may have important therapeutics and biotech implications. First, the mini-ribozyme has the simplest secondary structure with two stems and two internal loops, compared to all naturally occurring self-cleaving ribozymes. The previous minimum number of stems is three stems for hammerhead (33). Second, this mini-ribozyme after removing the stem-loop region only requires 17-nucleotide enzyme strand and 18-nucleotide substrate strand for its function. In other words, it can be easily modified as a trans-cleaving ribozyme, which has proven useful for bioengineering and RNA-targeting therapeutics (34). A recent intracellular selection based study has shown that hammerhead ribozyme can be engineered into a trans-acting ribozyme with consistent gene knockdown ability on different targeted mRNA either in prokaryotic or eukaryotic cells (34). In their work the most promising hammerhead variant contains 67 nt, with k_obs_ ~ 0.005 min^−1^ measured at a similar experimental condition (0.5mM Mg^2+^, pH 7.4) as our LINE-1-mini. The LINE-1 mini-ribozyme, the shortest one known, with higher cleavage rate than the most popular hammerhead ribozyme, may have an unprecedented advantage, compared to existing trans-cleaving ribozymes.

## Materials and Methods

All the oligonucleotides listed in Supplementary Table S4, except for the doped library, were purchased from IDT (Integrated DNA technologies) and Sigma-Aldrich. The doped mutant library Rz_LINE-1min_doped with a doping rate of 6%, was purchased from the Keck Oligo Synthesis Resource at Yale University. All the high-throughput sequencing experiments were performed on Illumina HiSeq X platform by Novogene Technology Co., Ltd.

### Deep mutational scanning experiments

The workflow for two deep mutational scanning experiments is illustrated in Supplementary Figure S8. The deep mutational scanning of the full-length LINE-1 (LINE-1-fl) ribozyme followed the same protocol of our previous work (16). Briefly, the mutant library was generated from 3-rounds of error-prone PCR, barcoded using primer Bar F and Bar R (Supplementary Table S4), and then diluted. After that the transcribed RNA of the barcoded mutant library was reverse transcribed with RT m13f adp1 and template switching oligo TSO (Supplementary Table S4). The total cDNA was used to generate the RNA-seq library with primer P5R1 adp1 and P7R2 adp2 (Supplementary Table S4). The DNA-seq libraries were generated by amplifying the ribozyme mutant library with primer P5R1_m13f and P7R2_t7p (Supplementary Table S4). Both DNA-seq and RNA-seq libraries were sequenced on an Illumina HiSeq X sequencer with 25% PhiX control by Novogene Technology Co., Ltd. We counted the numbers of reads of the cleaved and uncleaved portions of each variant by mapping their respective barcodes. The relative activity (RA) of each variant is calculated by using the equation *RA*(*var*) = *N*_*cleaved*_ (*var*)*N*_*total*_(*wt*)/*N*_*total*_(*var*)*N*_*cleaved*_ (*wt*).

The second round of deep mutational scanning experiment was applied to the minimal, contiguously functional region of LINE-1 ribozyme (LINE1-mini). Here, we used a doped mutation library instead of a library generated by error-prone PCR. The doped mutant library Rz_LINE-1_min_doped (Supplementary Table S4) was amplified by using primer T7prom and M13F (Supplementary Table S4). The PCR product was then gel-purified, quantified, and diluted. Approximately 5×10^5^ doped DNA molecules were amplified by T7prom and M13F primers (Supplementary Table S4) to produce enough DNA templates for in vitro transcription. The transcribed and purified RNAs (10 pmol) of the mutant library were mixed with 100 pmol rM13R_5desBio_3P (Supplementary Table S4), 2 μl 10X RtcB reaction buffer (New England Biolabs), 2 μl 1mM GTP, 2 μl 10mM MnCl_2_, 0.2 μl of Murine RNase Inhibitor (40 U/μl, New England Biolabs) and 1 μl RtcB RNA ligase (New England Biolabs) to a total volume of 20 μl. The reaction mixture was incubated at 37◦C for 1 h to allow higher cleavage ratio, then purified using RNA Clean & Concentrator-5 kit (Zymo Research). The purified products were mixed with 2 μl of 10 μl M RT_m13f_ adp1 (Supplementary Table S4) and 1 μl of 10mM dNTPs in a volume of 8 μl, and then heated to 65◦C for 5min and placed on ice. Reverse transcription was initiated by adding 4 μl of 5× ProtoScript II Buffer (New England Biolabs), 2 μl of 0.1 M DTT, 0.2 μl of Murine RNase Inhibitor (40 U/μl, New England Biolabs) and 1 μl ProtoScript II RT (200 U/μl, New England Biolabs) to a total volume of 20 μl. The reaction mixture was incubated at 42◦C for 1 h, and then purified by Sera-Mag Magnetic Streptavidin-coated particles (Thermo Scientific).

The purified products after streptavidin-based selection were amplified by PCR to construct the RNA-seq library with primer P7R2_m13r and P5R1_m13f (Supplementary Table S4). While the DNA template used for transcription was used as the template for constructing the DNA-seq library with primer P5R1 m13f and P7R2 t7p (Supplementary Table S4). The DNA-seq and RNA-seq libraries were also sequenced on an Illumina HiSeq X sequencer with 25% PhiX control by Novogene Technology Co., Ltd. Raw paired-end sequencing reads were filtered and then merged by using the bioinformatic tool PEAR (35) to generate the high-quality merged reads. We counted the read numbers of each variant in DNA-seq and RNA-seq by mapping the barcode. Then, we estimated the relative activity (RA’) of each variant, by using the equation *RA*′(*var*) = *N*_*RNA*_(*var*)*N*_*DNA*_(*wt*)/*N*_*DNA*_(*var*)*N*_*RNA*_(*wt*).

This relative activity (RA’) measure is different from the relative activity (RA) when both cleaved and uncleaved sequences were obtained in RNAseq in the first-round deep mutational scanning. However, it should be a good estimate of RA because a mutant with a higher activity (cleavage rate) should have a higher possibility to be captured by RtcB ligation. We confirmed RA’ as a good estimate of RA by comparing shared single mutations from LINE-1-fl and LINE-1-mini and obtained a strong Pearson correlation coefficient of 0.66 (Spearman correlation coefficient of 0.769). Moreover, both RA and RA’ (18-30 nt) show that the central region of LINE-1-mini is insensitive to mutations (Supplementary Figure S9). Moreover, secondary structures derived from RA’ and RA are consistent with each other (see below).

### Covariation-induced deviation of activity (CODA) analysis

We used CODA (16) to analyze the RA of LINE-1-fl and RA’ of LINE-1-mini. For Monte Carlo (MC) simulated annealing, we used the same weighting factor of 2 for both datasets as employed previously (16).

### PAGE-based cleavage assays

L1-S1-FAM (Supplementary Table S4) with 5′ 6-FAM as the fluorophore was synthesized and purified. In a 10 μl reaction system, 10 pmol L1-S1-FAM was mixed with 1 μl 0.2 M Tris-HCl and 0.5 μl 20 mM metal ion stock solution unless otherwise noted. 12 pmol L1-E2 (Supplementary Table S4) was added to initiate the reaction. Reactions were incubated at 37 ◦C for 1 h, and then stopped by adding an equal volume of stop solution (95% Formamide, 0.02% SDS, 0.02% bromophenol blue, 0.01% Xylene Cyanol, 1mM EDTA). The reaction products were then separated by denaturing (8 M urea) 15% PAGE, and imaged by using the Bio-Rad ChemiDoc XRS+ System.

Ribozyme cleavage assays for determining k_obs_ values were performed using the same bimolecular construct and experimental condition with 1mM Mg^2+^. Cleavage reactions were terminated using a stop solution containing 90% formamide, 50 mM EDTA, 0.05% xylene cyanol and 0.05% bromophenol blue at different time points. The fraction of 5′ 6-FAM labelled substrate RNA cleaved over time was quantified after separation by denaturing PAGE as described above. First order rate constants were determined by nonlinear curve fitting.

Cytosine C10 was substituted by deoxycytosine in L1-S1-dC10 (Supplementary Table S4) during synthesis. In a 10 μl reaction system, 10 pmol L1-S1-FAM or L1-S1-dC10 was mixed with 1 μl 0.2 M Tris-HCl, 0.5 μl 20 mM Mg^2+^ stock solution and 12 pmol L1-E2 (Supplementary Table S4). Reaction mixtures were incubated at 37 ◦C for 1 h, and then stopped for PAGE separation.

All the other mutants were generated by PCR amplification of two corresponding primers. Then the RNAs of these mutants were produced from 5 hours of in vitro transcription. The active ribozymes were co-transcriptionally cleaved, and the reaction mixtures were stopped and separated by denaturing (8M urea) 15% PAGE and visualized by SYBR Gold (Thermo Fisher) staining.

### Homology search of LINE-1-mini

Sequence-based search of LINE-1-mini. The Basic Local Alignment Search Tool (BLAST-N) (25) was used to search homologs of LINE-1-mini ribozyme in the NCBI Refseq Representative genomes Database (https://ftp.ncbi.nlm.nih.gov/blast/db/) with E-value setting to 0.001. This database covers 1000+ representative eukaryotic genomes, 5700+ representative prokaryotic genomes, 9000+ representative virus genomes and 46 representative viroid genomes in total. We used blastdbcmd from BLAST and in-house scripts to fill the gaps at both two ends for alignments.

Secondary structure-based search of LINE-1-mini. All representative primate genome assemblies in fasta format and the corresponding RepeatMasker (http://www.repeatmasker.org) annotation files were downloaded from the UCSC genomic website (http://genome.ucsc.edu). We used cmbuild from Infernal (36) to build the covariance model (26) of LINE-1-mini ribozyme with the wild type sequence and secondary structure inferred from deep mutational scanning experiment as input. Here, we only consider Watson-Crick pairs (WC-pairs), which form one stem of 5 base pairs and another stem of 6 base pairs. Afterwards, we used the covariance model to search against the latest release of different representative primate genome assemblies by using cmsearch from Infernal. The hits with E-value <1.0 were then used for downstream analysis.

## Data Availability

Illumina sequencing data for the LINE-1-fl and LINE-1-mini were submitted to the NCBI Sequence Read Archive (SRA) under SRA accession number PRJNA662002 (https://www.ncbi.nlm.nih.gov/sra/PRJNA662002).

## Code availability

All custom scripts needed to repeat the analyses are available at https://github.com/zh3zh/CODA2.

## Acknowledgment

We would like to thank Jaswinder Singh for helping us to predict RNA secondary structure by SPOT-RNA. Z.Z. gratefully acknowledges the award of Griffith University and University of Chinese Academy of Sciences (GU-UCAS Joint Doctoral Program) Postgraduate Scholarships. This work was supported in part by Australia Research Council DP180102060 and DP210101875 and by National Health and Medical Research Council (1121629) of Australia to Y.Z. We would like to acknowledge the use of the High-Performance Computing Cluster ‘Gowonda’ to complete this research. This research/project has also been undertaken with the aid of the research cloud resources provided by the Queensland Cyber Infrastructure Foundation (QCIF) and Australian Research Data Commons (ARDC). We also gratefully acknowledge the support of NVIDIA Corporation with the donation of the Titan V GPU used for this research.

## Author Contribution

JZ conceived of the study. JZ and JW designed experimental studies and wrote the manuscript; ZZ and JZ conducted experimental studies and wrote the manuscript; PX developed computational methods. ZZ performed computational and bioinformatics analysis. YZ participated in the initial design, assisted in data analysis, and drafted the manuscript; and all authors read, contributed to the discussion, and approved the final manuscript.

