## Supplementary materials for "Resurrecting self-cleaving mini-ribozymes from 40-million-year-old LINE-1 elements in human genome"

Table S1. The number of mutants and the coverage of single and double mutations of LINE-1 ribozyme in two deep mutational scanning experiments for full-length (LINE-1-fl) and minimal functional regions (LINE-1-mini), respectively.

| Mutations | Single mutation | | | Double mutation | | |
|---|---|---|---|---|---|---|
| | Number | Total | Coverage | Number | Total | Coverage |
| LINE-1-fl | 429 | 438 | 97.9% | 17671 | 95265 | 18.5% |
| LINE-1-mini | 137 | 138 | 99.3% | 9310 | 9315 | 99.9% |

Table S2. Summary of two deep sequencing results for full-length (LINE-1-fl) and minimal functional regions (LINE-1-mini), respectively.

| | Number of DNA-seq reads | Number of Barcodes | Number of mutants | Number of RNA-seq reads | Cleaved | Uncleaved |
|---|---|---|---|---|---|---|
| LINE-1-fl | 89053476 | 168221 | 70962 | 59208161 | 1982615 | 39512450 |
| LINE-1-mini | 66952586 | 373301 | 162752 | 69904460 | 69904460 | NA |

Table S3. Secondary-structure-based search result of LINE-1-mini in UCSC representative genome assemblies by using cmsearch

| Species | Genome assembly ID (UCSC) | Number of hits (E-value<=1.0) | Number of hits matched to LINE-1 region |
|---|---|---|---|
| Human | hg38 | 1142 | 1142 |
| Chimpanzee | panTro6 | 1107 | 1107 |
| Bonobo | panPan2 | 1052 | 1052 |
| Gorilla | gorGor5 | 1087 | 1087 |
| Orangutan | ponAbe3 | 1100 | 1100 |
| Gibbon | nomLeu3 | 956 | 955* |
| Green monkey | chlSab2 | 973 | 973 |
| Crab-eating macaque | macFas5 | 980 | 978 |
| Rhesus macaque | rheMac10 | 992 | 991 |
| Baboon (anubis) | papAnu4 | 934 | 932 |
| Baboon (hamadryas) | papHam1 | 920 | 918 |
| Proboscis monkey | nasLar1 | 731 | 731 |
| Golden snub-nosed monkey | rhiRox1 | 971 | 969 |
| Marmoset | calJac3 | 601 | 601 |
| Squirrel monkey | saiBol1 | 531 | 531 |
| Tarsier | tarSyr2 | 0 | 0 |
| Mouse lemur | micMur2 | 1 | 0 |
| Bushbaby | otoGar3 | 0 | 0 |

*there is one hit which matches to the LINE-1 element in a reverse direction

Table S4. Oligonucleotides used in this study.

| Name | Sequence (5’ - 3’) | Notes |
|---|---|---|
| Rz_LINE-1_wt | TAATACGACTCACTATAGGGATACGTAGGTGGTTTTCCCCTGATAGTGCTAAGGGGGCCTGGAAGTTTGGACTGGGCAGAGCTCAACACAGCGTGGCAAAGTGGCTGTGGCTAGACTGCCTCTCTGGATTCCTCGTCACTCAGCAGGGCATCTCTGAGAGAAATGCCATTTGAGTTACTCAGGATTACTGGCCGTCGTTTTAC | DNA template. |

| | | |
|---|---|---|
| Rz_LINE-1_min_doped | TAATACGACTCACTATAGGGGCAGAGCTCAACACAGCGTGGCAAAGTGGCTGTGGCTAGACTGCCAANNNNNNNNNNNNNNNNNNNACTGGCCGTCGTTTTAC | Doped library. |
| M13F | GTAAAACGACGGCCAGT | PCR primer. |
| T7prom | TAATACGACTCACTATAGGG | PCR primer. |
| RT_m13f_adp1 | CCCTACACGACGCTCTTCCGATCTGTAAAACGACGGCCAGT | Reverse transcription primer. |
| TSO | CTCGGCATTCCTGCTGAACCGCTCTTCCGATCTrGrGrG | Template switching oligo. |
| rM13R_5desBio_3P | /5deSBioTEG/rCrArCrArGrGrArArArCrArGrCrUrArUrGrArCrC/3Phos/ | Linker for RtcB ligation. |
| Bar_F | TAATACGACTCACTATAGGGA | PCR primer for adding barcode. |
| Bar_R | GTAAAACGACGGCCAGTNNNNNNNNNNNNNNNNAATCCTGAGTAACTCAAAT | PCR primer for adding barcode. |
| P5R1_m13f | AATGATACGGCGACCACCGAGATCTACACTCTTTCCCTACACGACGCTCTTCCGATCTGTAAAACGACGGCCAGT | PCR primer for DNA-seq library preparation. |
| P7R2_t7p | CAAGCAGAAGACGGCATACGAGATCGGTCTCGGCATTCCTGCTGAACCGCTCTTCCGATCTTAATACGACTCACTATAGG | PCR primer for DNA-seq library preparation. |
| P5R1_adp1 | AATGATACGGCGACCACCGAGATCTACACTCTTTCCCTACACGACGCTCTTC | PCR primer for RNA-seq library preparation. |
| P7R2_adp2 | CAAGCAGAAGACGGCATACGAGATCGGTCTCGGCATTCCTGCTGAAC | PCR primer for RNA-seq library preparation. |
| P7R2_m13r | CAAGCAGAAGACGGCATACGAGATCGGTCTCGGCATTCCTGCTGAACCGCTCTTCCGATCTCAGGAAACAGCTATGACC | PCR primer for RNA-seq library preparation. |
| L1-FAM-S1 | /56-FAMK/rGrGrCrArGrArGrCrUrCrArArCrArCrArGrC | Oligo for PAGE-based activity assay. |
| L1-E2 | rGrCrUrGrUrGrGrCrUrArGrArCrUrGrCrC | Oligo for PAGE-based activity assay. |
| L1-S1-dC10 | rGrGrCrArGrArGrCrUCrArArCrArCrArGrC | Oligo for PAGE-based activity assay. |
| PA7_for | TAATACGACTCACTATAGGGCAGAGCCCACCGCAGCT | PCR primer for PAGE-based activity assay. |
| PA7_rev | GGCAGTCTGGCTACAGCGGCTTTGCTGAGCTGCGGTGG | PCR primer for PAGE-based activity assay. |
| P3_for | TAATACGACTCACTATAGGGCAGAGCCCACCACAGCT | PCR primer for PAGE-based activity assay. |

| | | |
|---|---|---|
| P3_rev | GGCAGTCTGGCTACAGCGGCTTTGCTGAGCTGTGGTGG | PCR primer for PAGE-based activity assay. |
| PA8_for | TAATACGACTCACTATAGGGCGGAACTCACCACAGCG | PCR primer for PAGE-based activity assay. |
| PA8_rev | GGCAGTCTGGCCACAGCCGCTTTGCCGCGCTGTGGTGA | PCR primer for PAGE-based activity assay. |
| PA10_for | TAATACGACTCACTATAGGGCAGAATTCACCACAGCG | PCR primer for PAGE-based activity assay. |
| PA10_rev | AGCAGTCTGGCCACAGCCGCTTTGCTGCGCTGTGGTGA | PCR primer for PAGE-based activity assay. |
| P2_for | TAATACGACTCACTATAGGGCAGAGCCCACCACAGCA | PCR primer for PAGE-based activity assay. |
| P2_rev | GGCAGTCTGGCTACAGTGGTTTTGCCGTGCTGTGGTGG | PCR primer for PAGE-based activity assay. |
| L1_wt_for | TAATACGACTCACTATAGGGCAGAGCTCAACACAGCG | PCR primer for PAGE-based activity assay. |
| L1_wt_rev | GGCAGTCTAGCCACAGCCACTTTGCCACGCTGTGTTGA | PCR primer for PAGE-based activity assay. |
| L1_A11G_for | TAATACGACTCACTATAGGGCAGAGCTCGACACAGCG | PCR primer for PAGE-based activity assay. |
| L1_A11G_rev | GGCAGTCTAGCCACAGCCACTTTGCCACGCTGTGTCGA | PCR primer for PAGE-based activity assay. |
| L1_G7A,C8U_for | TAATACGACTCACTATAGGGCAGAATTCAACACAGCG | PCR primer for PAGE-based activity assay. |
| L1_T9C_for | TAATACGACTCACTATAGGGCAGAGCCCAACACAGCG | PCR primer for PAGE-based activity assay. |
| L1_T9C_rev | GGCAGTCTAGCCACAGCCACTTTGCCACGCTGTGTTGG | PCR primer for PAGE-based activity assay. |
| L1_A12C_for | TAATACGACTCACTATAGGGCAGAGCTCACCACAGCG | PCR primer for PAGE-based activity assay. |
| L1_A12C_rev | GGCAGTCTAGCCACAGCCACTTTGCCACGCTGTGGTGA | PCR primer for PAGE-based activity assay. |
| L1_G35A_rev | GGCAGTCTAGCTACAGCCACTTTGCCACGCTGTGTTGA | PCR primer for PAGE-based activity assay. |
| L1_T38C_rev | GGCAGTCTGGCCACAGCCACTTTGCCACGCTGTGTTGA | PCR primer for PAGE-based activity assay. |

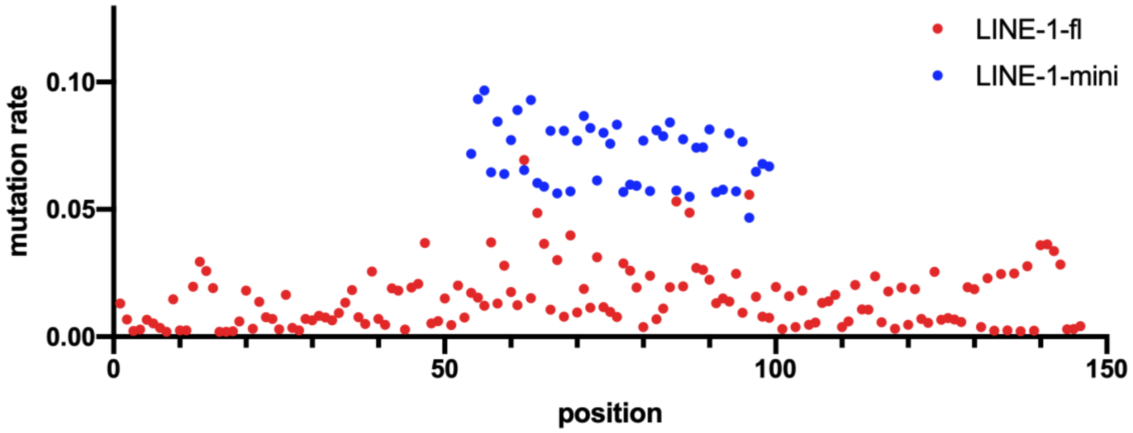


Figure S1. Mutation rates of two deep mutational scanning results at each nucleotide position of LINE-1 ribozyme.

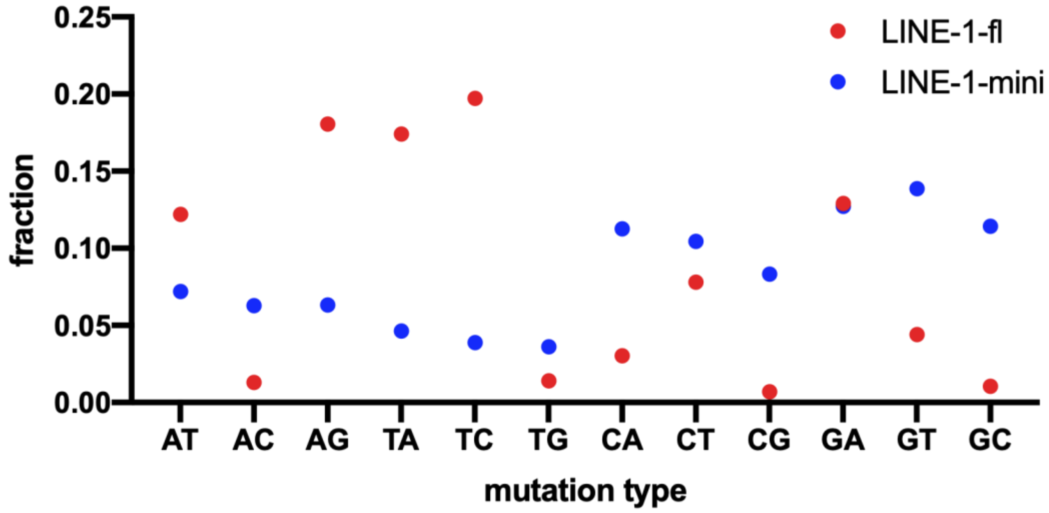


Figure S2. Distribution of different mutation types in deep sequencing result.

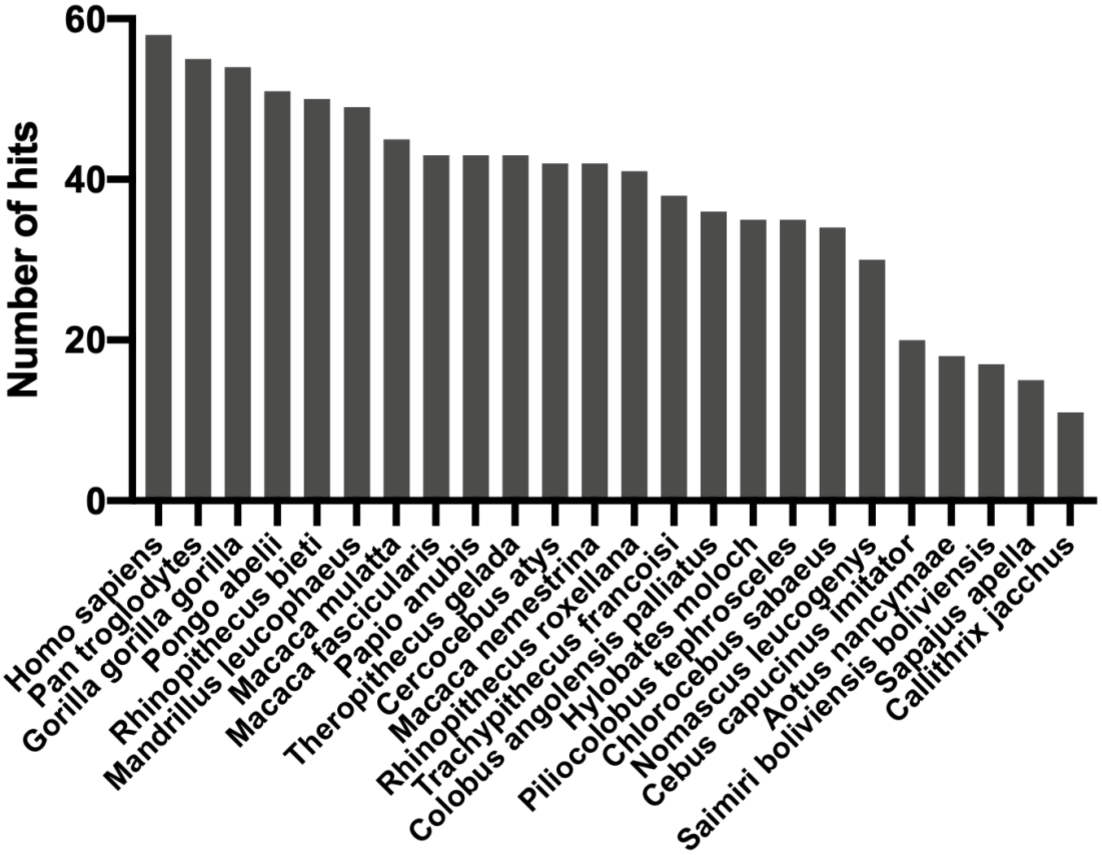


Figure S3. Number of LINE-1-mini homologs found in each species by using BLAST-N.

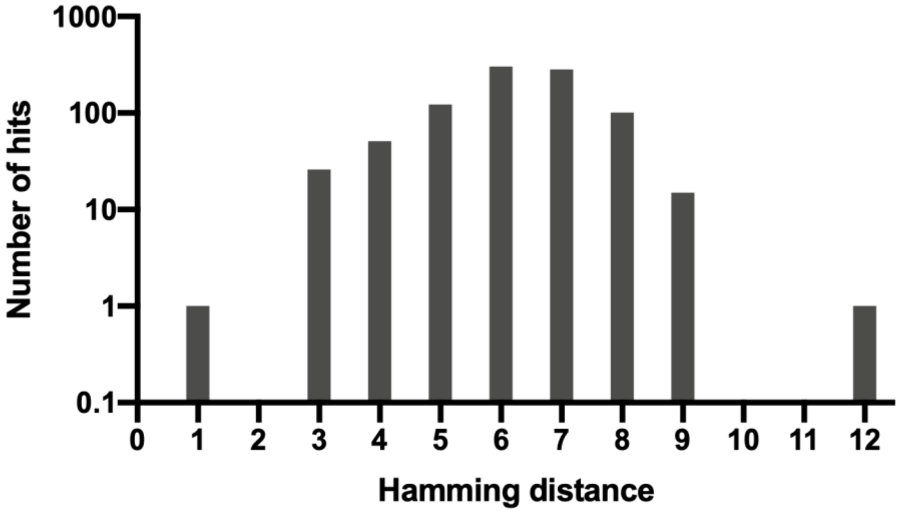


Figure S4. Hamming distance distribution of LINE-1-mini homologs found by using BLAST-N.

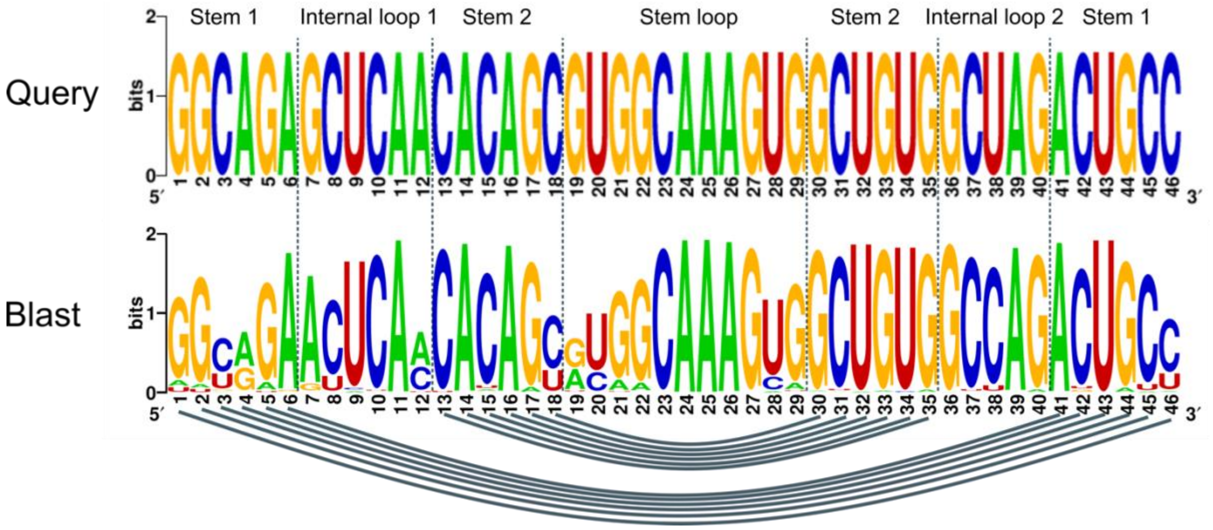


Figure S5. Nucleotide composition of LINE-1-mini homologs found by using BLAST-N.

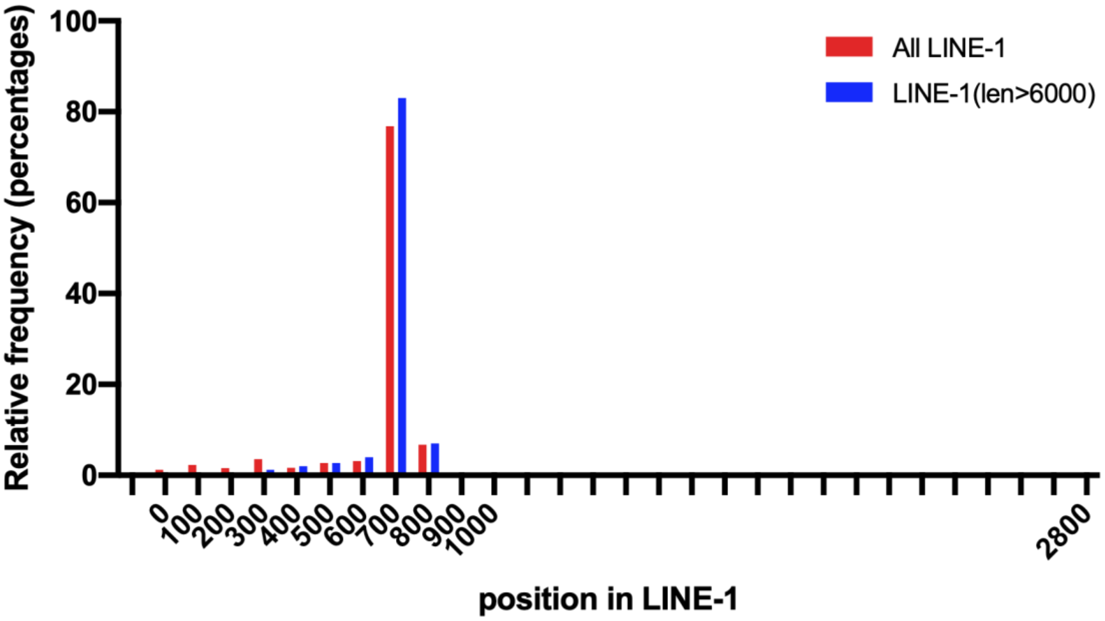


Figure S6. The frequency distribution of the position of LINE-1-mini homologs inside LINE-1 retrotransposons.

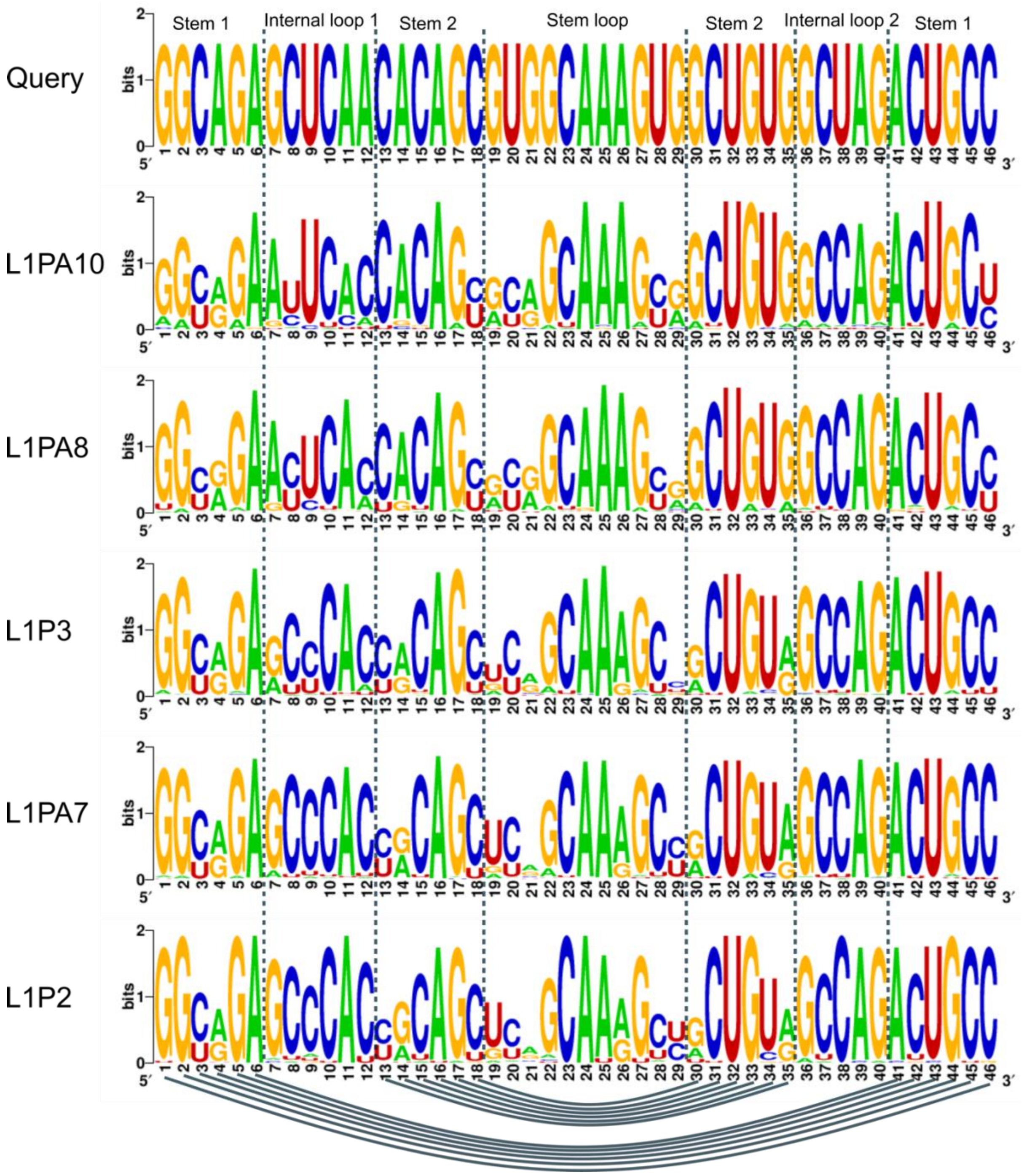


Figure S7. The comparison of nucleotide composition in the 5 most abundant subfamilies of LINE-1.

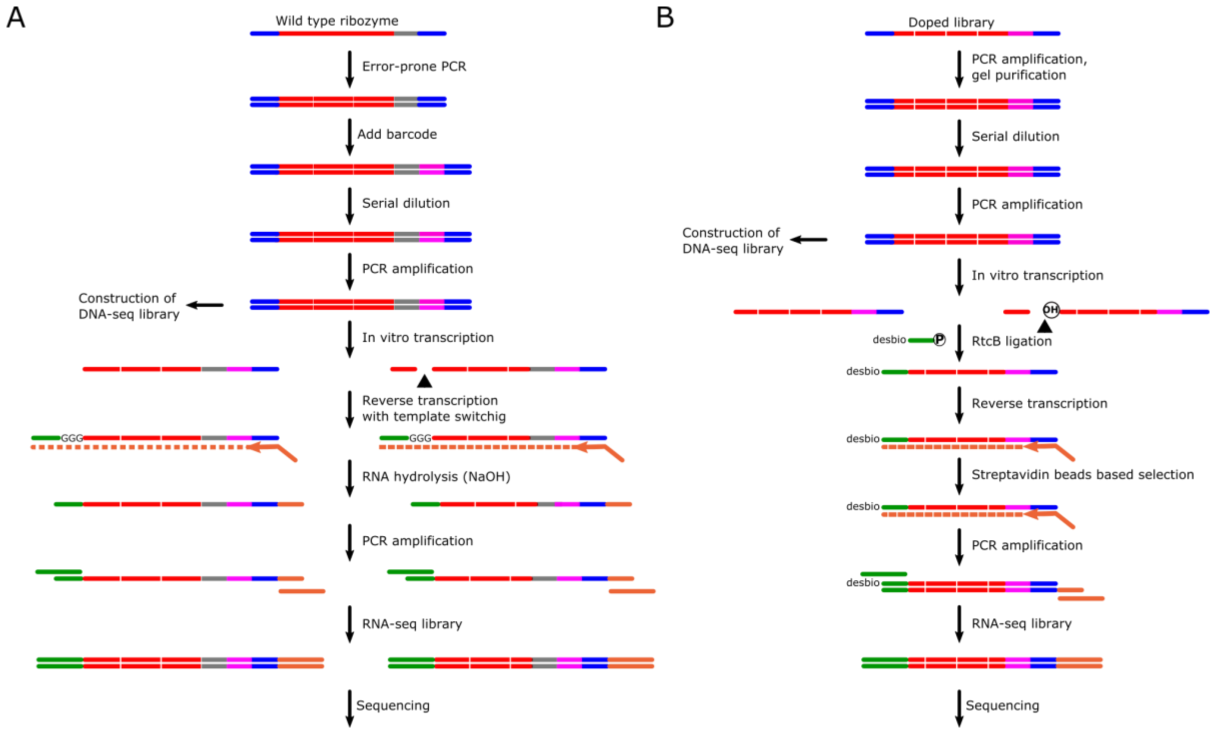


Figure S8. Experimental pipeline of two deep mutational scanning experiments. (A) The mutation library of full-length LINE-1 ribozyme was generated by error-prone PCR. The RNA-seq library was constructed by using the cDNA synthesized by reverse transcription and template switching. (B) The chemically synthesized doped library was used to construct the mutation library of the minimal, contiguous, functional region of LINE-1 ribozyme. RtcB ligase was used to captured 3′ fragment after cleavage, then the ligated product was used to construct the RNA-seq library.

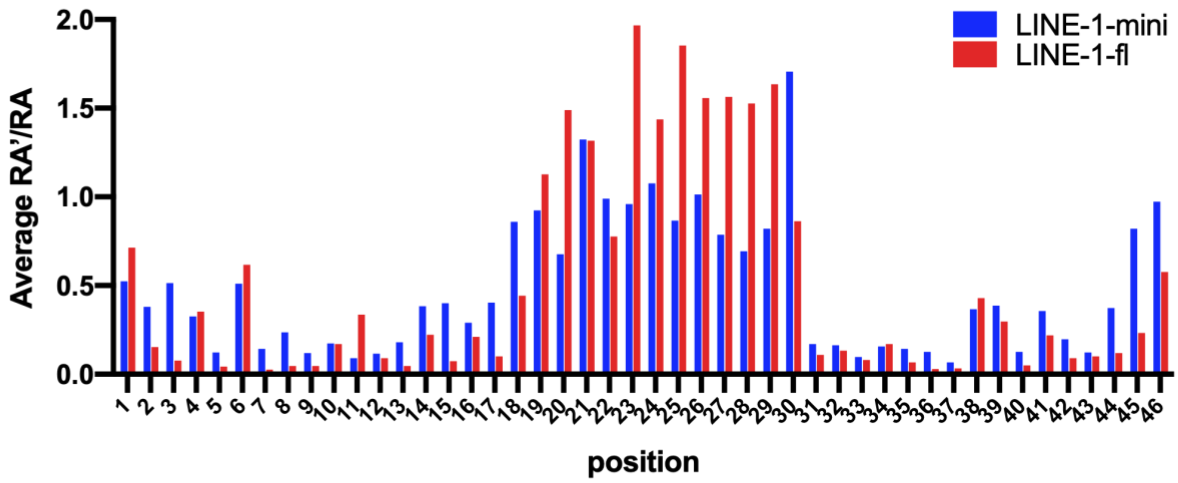


Figure S9. Single mutation-based average RA'/RA distribution of LINE-1-mini.
